# According to a large citizen science dataset, UK farmland bird indicator species and other bird groups, are more likely to be found on farms with shoots than on otherwise similar farms

**DOI:** 10.64898/2026.10.26.760633

**Authors:** R.B. Sage, L. Costick, A.M. Lole, E. Raynor, J.R. Egan, R.A.H. Draycott

## Abstract

On average around 1500 farms across lowland UK take part in the GWCT Big Farmland Bird Count in spring each year. Alongside bird data based on a 30-minute survey of 2ha of farmland, participants are asked a group of other questions including, for the last three years only, whether there is a gamebird shoot or not on the farm. The dataset is created from online forms and is generally of low quality including missing information and suspect bird ID. We suggest this does not cause bias in our investigation. The dataset does not include a site identifier, so we have analysed data year by year, and the caveats around that in interpretation are discussed. The standard 2ha survey patch on each farm is not selected randomly; larger farms might have more choice and shoots tended to be on larger farms. Removing the effect of farm size in analysis, as well as other factors including the use of Agri Environment Scheme (AES) options, our bird data suggest that farms with a shoot, have on average around 10% greater bird species richness than otherwise similar farms. This includes the UK Farmland Bird Index (FBI) species group and Red and Amber listed species. Raptor species were 20% more commonly encountered on shoots than non-shoots. Previous studies have shown several potential drivers of this, and we suggest the provision of game crops on most shoots that release gamebirds is a contributor. We estimate that game management, which includes the planting of game crops, is undertaken on around 20% of lowland UK farmland. Using this figure, according to our bird results, overall and FBI species richness would be about 2% lower across all of lowland UK farmland without game management. This does not include any benefit to birds associated with the use of AES options on shoots. Bearing in mind the limitations of the data and the analysis, our investigation suggests the privately funded contribution made by game management will be contributing to the maintenance of farmland bird populations on farms and, potentially, reducing the national FBI rate of decline.

## Introduction

Many farmland birds have suffered population declines across Europe since the 1970s (Gregory et al. 2004). There are a variety of demographic and environmental causes that have been identified (Newton, 2004) but broadly the main driver for many seed-eating farmland species is reduced overwinter survival caused by lack of food resource. For a group of non-passerine species (especially waders) and some passerines, population growth is often limited by productivity. Changes to within-field plant and invertebrate community has been linked primarily but not exclusively to the use of pesticides (Potts et al. 2010; Ewald et al. 2024).

These insights led to the development of a range of apparently focussed conservation interventions, including seed-rich and insect-rich foraging habitats and fallow plots (Wilson et al. 2009; Winspear et al. 2010). In the decade or so since the deployment of these interventions within Agricultural Environmental Schemes (AES) in the UK and elsewhere in Europe, there have been some successes, especially when an intervention has been targeted on a particular bird or problem (Prince et al. 2012; Redhead et al, 2022). The overall picture however is of continued declines in many farmland specialist species and to a lesser extent in generalist species using farmland habitats (Sharps et al. 2023).

The documentation of this in the UK is shown in the breeding Farmland Bird Indicator (FBI) (Eaton et al. 2026). This index has now declined by over 60% since 1970. Most of the decline occurred in the late 1970s and 1980s followed by a reduced rate of decline from around 1990. However, in the last few years the index decline has accelerated resulting in a 5-year reduction of around 10% up to 2026.

The main mechanism for the deployment of conservation interventions on farmland other than through AES options is through privately funded actions by farmers and other land managers on their own land. Within this broad area of often ‘under-valued’ conservation work there is game management. There are several reasonably well documented mechanisms by which management of farmland for released (and wild) gamebirds might improve over-winter survival, spring densities and potentially breeding success of wild birds using that farmland (reviewed in Sage et al. 2020; Madden & Sage 2020). These include the use of game crops (Sage et al. 2005), food resources (Siriwardena et al. 2006), better hedgerow and wood edge habitats (Draycott et al. 2012), and predator control (Roos et al. 2018).

On shoots that release gamebirds there are also possible effects of high densities of pheasants and/or red-legged partridges on a site that may negatively affect farmland birds. Wood edge and hedgerow habitats can be degraded (Sage et al. 2020).

There are possible zoonotic disease issues that might affect some birds (Madden & Sage 2020). It has been shown that where birds congregate, either at release pens or potentially at feeding sites, soils and flora can be degraded and invertebrate communities altered (Sage et al. 2005; Sage 2026).

Clearly game management is not preventing national declines in farmland birds but the question remains whether declines would be faster and steeper, or possibly less severe, without it? The studies on the various consequences of releasing gamebirds, reviewed in Sage et al. (2020) and explored in the Discussion in this paper, usually involve dedicated field studies at a sample of sites, sometimes only two or three, with many others involving upwards of ten plus sites.

GWCT now has a dataset on bird use on a large sample of farms based on an online questionnaire called the Big Farmland Bird Count (BFBC). While the questionnaire is not focussed on game related matters, in recent years the information collected includes whether there is a shoot on the farm or not. The question considered by our investigation of the BFBC dataset in this paper is, can the findings of the focussed studies on bird use of habitat patches on game estates be extrapolated to a very large sample of game managed farmland sites throughout the country? And if so, to what extent might game management be influencing farmland bird abundance on lowland agricultural land in the UK as a whole.

## Methods

### Surveys

The Big Farmland Bird Count (BFBC) is a citizen science initiative which aims to measure how farmland birds are faring and to highlight the conservation role of farmers, rather than for scientific investigation. It uses a simple 30-minute survey carried out on farms across Great Britain each February by farmers, land managers and volunteers. The first count in 2014 involved around 500 landowners, and by 2021 it had grown to a record 2,500 count returns. The count is done during a two-week window, scheduled for February, as it is designed to capture a snapshot of farmland bird populations during the winter months. This is when many species are most visible and when agricultural land management practices have a direct impact on their survival. February is widely referred to as the ‘hungry-gap’, (Stoate, 2012) when nutritional resources are at their lowest for farmland birds and interventions by farmers such as food plots or late winter supplementary feeding can benefit farmland birds (BES 2013). It is also a time of year when many farmers have some spare time in their busy daily schedules, relative to other times of year, e.g. harvest.

Counts are recorded in the field on paper and subsequently uploaded directly to a digital platform within the Big Farmland Bird Count website, or, since 2026, directly via a mobile phone app. Counts are undertaken in 2ha plots, chosen by the farmer and the maximum number of each species seen at any one time is recorded (to reduce double-counting) during a 30-minute period. In addition to species and numbers seen, other factors are recorded, including location, date, time of day, weather conditions, habitat and cropping and whether farms were in agri-environment schemes (AES). In the last three years, 2024-2026 participants have also been asked whether there is a shoot on the farm or not. This is the only game management related question in the survey.

### Analysis

To facilitate our simple scientific investigation, we first cleaned the bird abundance data, primarily looking for and removing or otherwise dealing with obvious errors. We then converted all bird counts into simple presence/absence data, intending to reduce reliance on inconsistent and potentially very inaccurate abundance estimates. Birds that did not appear in all three years were assumed to be absent from counts in those years. Using this presence/absence data, we created species richness variables, with species richness defined here as the number of unique species per count. Along with total species richness, we calculated richness of the following groups of birds:

- Farmland Bird Indicator species (Eaton et al., 2026);
- Birds of Conservation Concern - Amber and Red listed species (Stanbury et al., 2021);
- Three taxonomic groupings: finches and buntings; raptors; thrushes.

The three datasets were analysed separately as counts from the same farm/estate could not be reliably linked between years.

All statistical analyses were conducted using R Statistical Software (R Core Team, 2026) in RStudio (Posit team, 2026), using the following packages: tidyr (Wickham et al., 2014), DHARMa (Hartig, 2016), and MASS (Venables et al., 2002).

We used shoot status (yes/no) as the focal explanatory variable. Poisson GLMs were initially fitted; however negative binomial models using a logarithmic link function were ultimately chosen to account for overdispersion in the richness data. We first created unadjusted models which estimated the relationship between species richness and shoot status only. We then selected covariates based on ecological relevance, with consideration of data completeness to create a model most representative of the data. Farm size was selected as a covariate due to the species-area relationship which shows that the number of species present increases with area (Connor and McCoy, 1979). While the bird counts were carried out over a 2ha patch of land, farm size still has the potential to influence species richness as the 2ha patch was not randomly selected. Farm size values of zero or extremely implausible values were treated as missing, then the variable was log transformed to accommodate its right-skewed distribution. Region was also selected as a covariate to account for geographical variation in species distributions. AES participation and Cluster Farm membership were selected as covariates as they may be associated with environmental management practices that influence richness and differ between shoot and non-shoot farms, so could therefore be a potential confounder of the shoot status ∼ species richness association. Cluster Farms are voluntary associations between bordering farms which undertake modest conservation work over a wide area (Nichols et al. 2025).

The adjusted models used a complete-case dataset, only observations with complete data for shoot status and all covariates were used.

Unadjusted model formula:

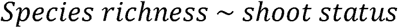

Adjusted model formula:

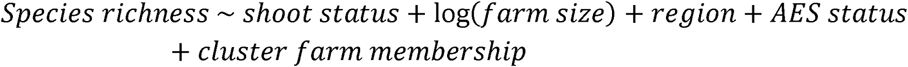

Results were expressed using the incidence rate ratio (IRR) and 95% confidence intervals. The reference category for shoot status was the non-shoot farms; the IRR compares shoot with non-shoot farms.

Model adequacy was assessed using simulated residual diagnostics generated with the DHARMa package, which included tests for uniformity, dispersion, and zero inflation. Potentially influential observations were identified using Cook’s distance.

Sensitivity analyses were conducted on the adjusted FBI richness models by restricting the dataset to observations from England. Additionally, the 2025 FBI richness model was refitted after excluding the ten observations with the highest Cook’s distances to assess the influence of these observations on the estimated association between shoot status and species richness.

## Results

### Dataset characteristics

The data contained a total of 4829 observations across 2023-2025. Participants reported that about 30% of observations were on arable farmland, 40 % on grass and the remainder were mixed. Of the total observations, 84.86% (n = 4098) contained valid information on shoot status. Of these, 3325 observations (68.85%) had complete data for shoot status and the covariates included in the adjusted models. Covariate summary statistics (excluding region) are shown in Table 1.

**Table 1.** Summary of covariate statistics excluding region. Percentages shown for AES participation and Cluster Farm membership are how many participants answered “yes” each year.

| Year | Shoot status | N | Median farm size (ha) | AES participation (%) | Cluster farm membership (%) |
| --- | --- | --- | --- | --- | --- |
| 2023 | Yes | 419 | 300 | 83.1 | 20.4 |
|  | No | 703 | 80 | 47.4 | 29.8 |
| 2024 | Yes | 464 | 300 | 77.8 | 21.6 |
|  | No | 745 | 89 | 53.2 | 29.7 |
| 2025 | Yes | 415 | 300 | 78.3 | 23.5 |
|  | No | 579 | 95.5 | 57.2 | 39.0 |

Median farm size was consistently greater for shoot farms (300ha in all three years) compared to non-shoot farms (80-95.5ha). AES participation was consistently higher in shoot farms (77.8-83.1%) compared to non-shoot farms (47.4-57.2%).

Observations were distributed across UK regions, with region included as a covariate for adjusted models.

### Shoot status and species richness

Overall, species richness was generally positively associated with shoot status after adjusting for farm size, region, AES participation, and Cluster Farm membership.

However, the magnitude and statistical significance of these associations varied between years and between bird groups (Table 2).

**Table 2.**
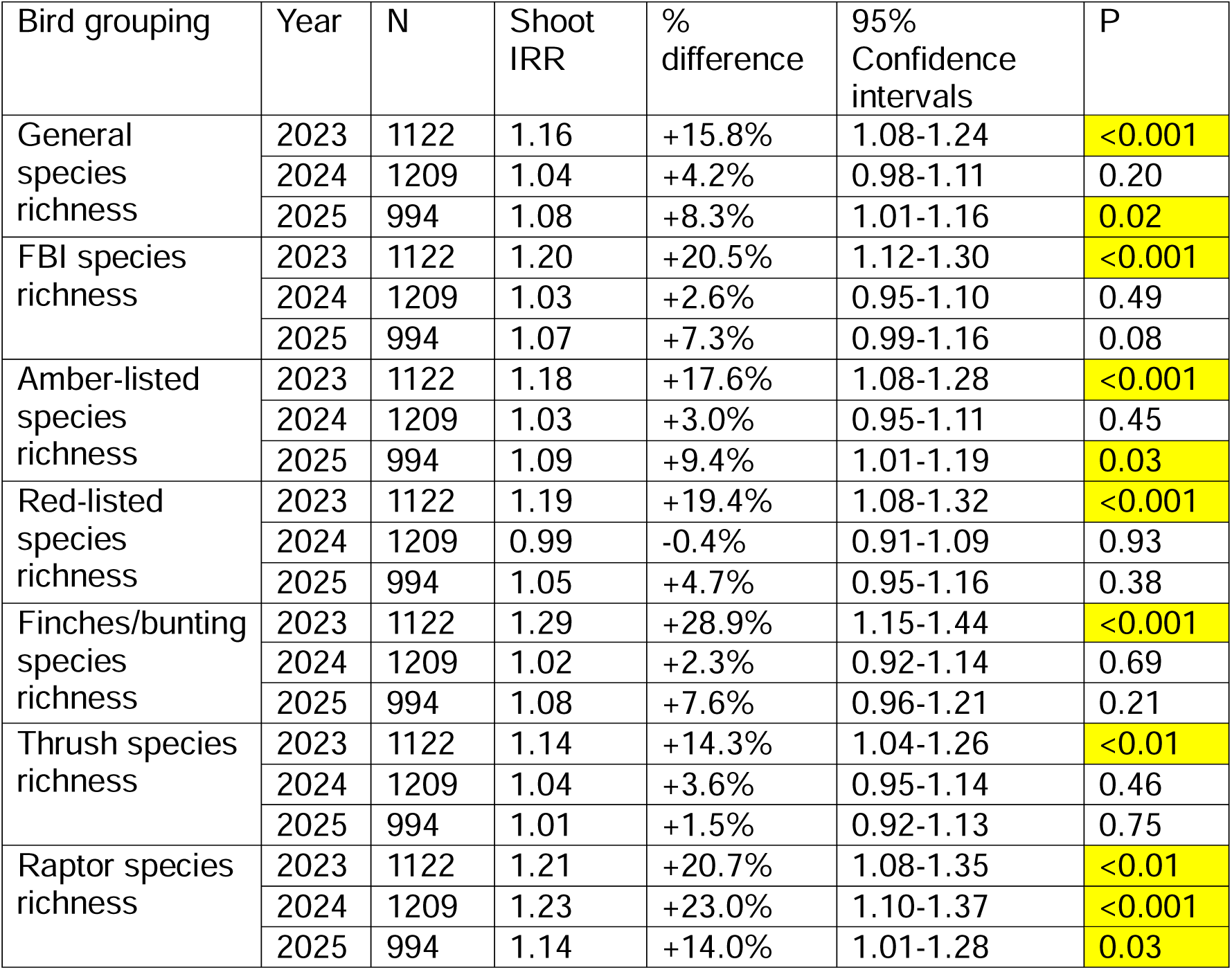
Adjusted model results for all bird group species richness for each year.

For overall species richness, shoot farms had an estimated 15.8% higher species richness (p < 0.001) in 2023, and an estimated 8.3% higher species richness (p = 0.02) in 2025 than non-shoot farms, after accounting for covariates. In 2024, estimated species richness was 4.2% higher on shoot farms; however, the association was not statistically significant (p = 0.20). Model adjusted mean species richness across the three years is presented in Figure 1.

**Figure 1.**
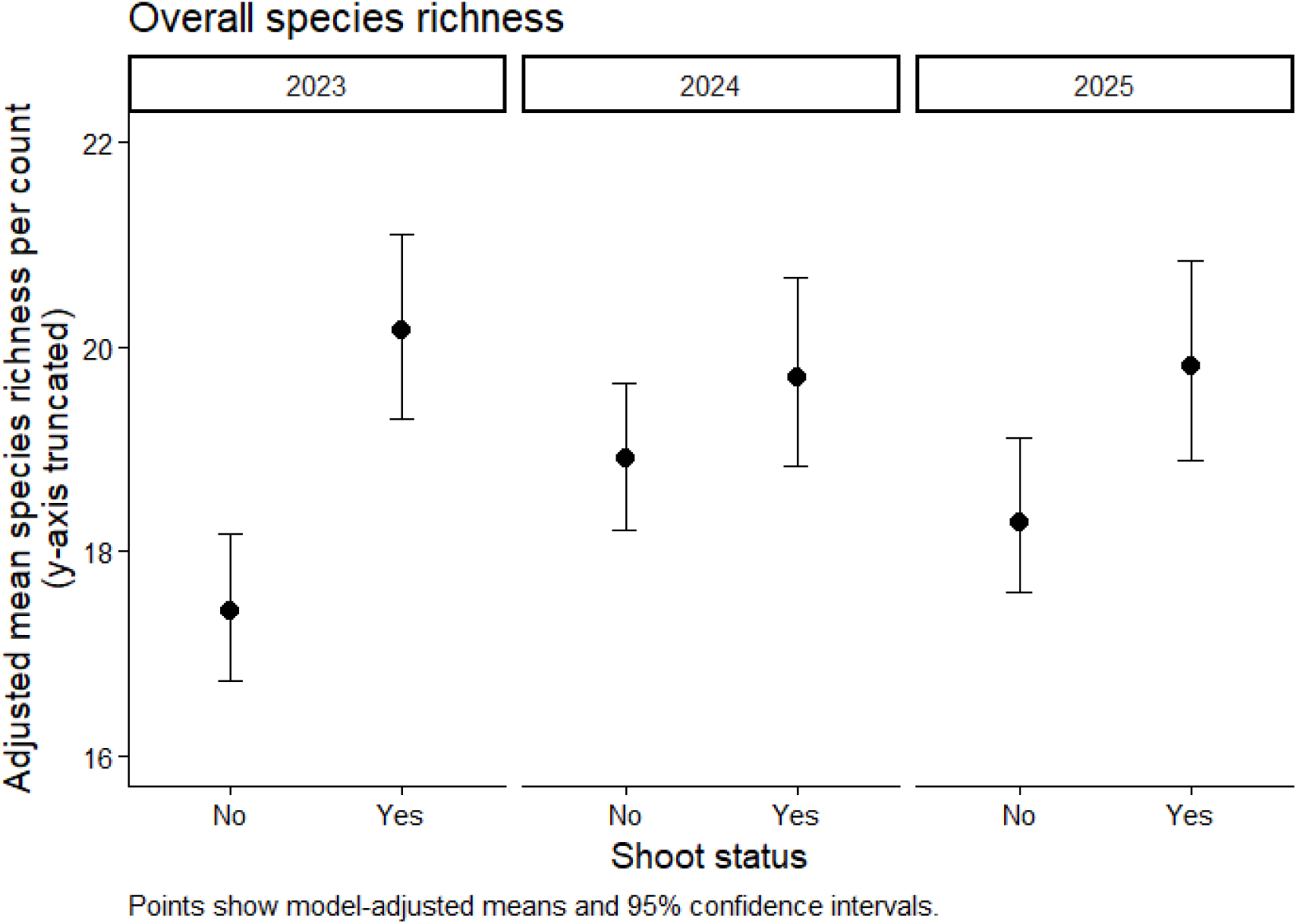
Model-adjusted mean overall bird species richness per count in shoot and non-shoot farms, 2023–2025. Points represent adjusted means and error bars indicate 95% confidence intervals.

A similar pattern was observed for FBI species richness, with the strongest positive association in 2023, where estimated richness was 20.5% higher on shoot farms (p < 0.001). Smaller positive associations were estimated in 2024 (+2.6%, p = 0.49) and 2025 (+7.3%, p = 0.08), although neither was statistically significant (Figure 2).

**Figure 2.**
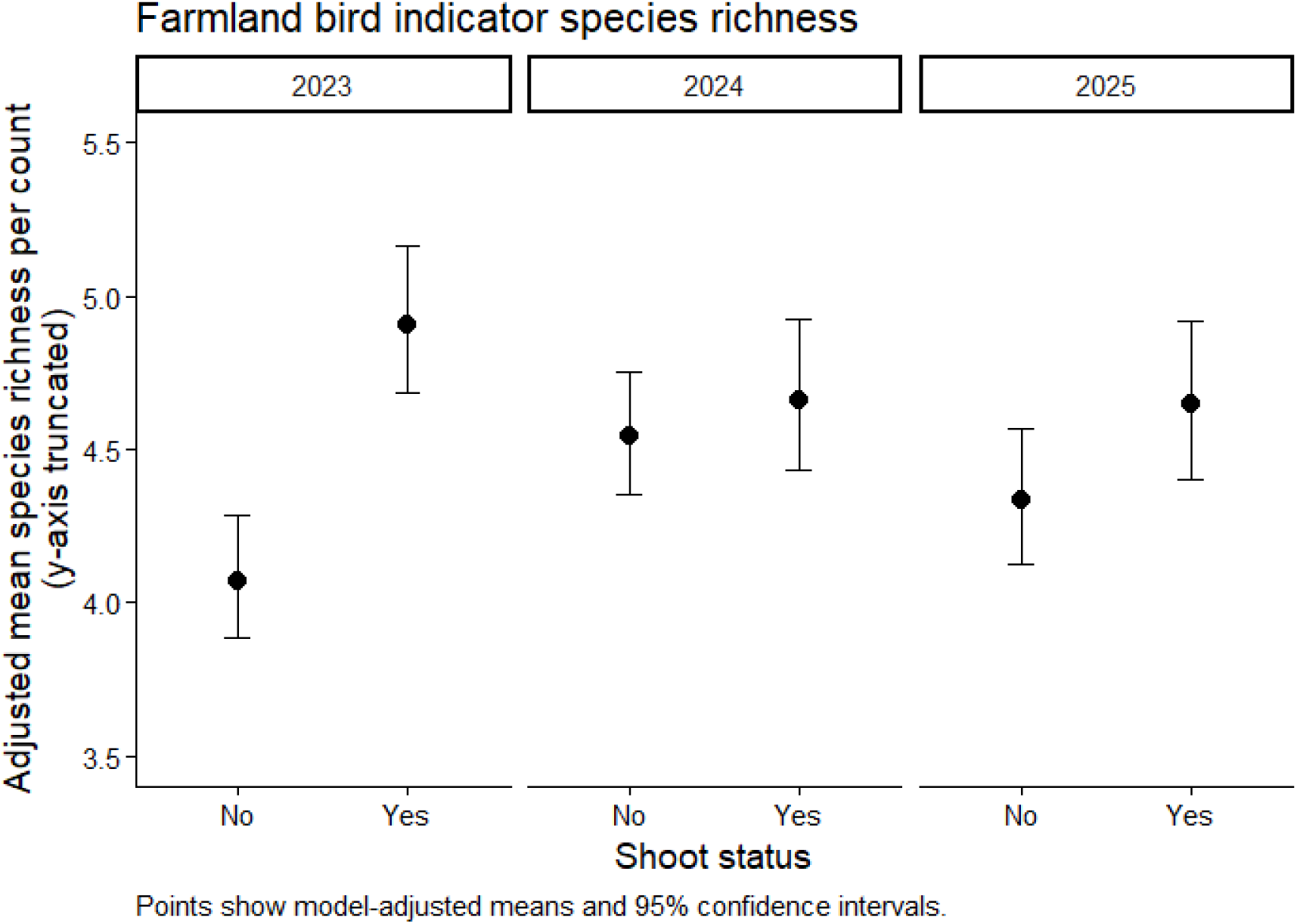
Model-adjusted mean FBI species richness per count in shoot and non-shoot farms, 2023–2025. Points represent adjusted means and error bars indicate 95% confidence intervals.

Positive associations were observed for both Amber- and Red-listed species richness in 2023, with estimated richness 17.6% and 19.4% higher on shoot farms respectively (both p < 0.001). In 2024 associations were small and not statistically significant for either group. In 2025, the estimated association was positive for both groups, although only Amber-listed species richness showed a statistically significant association (p = 0.03; Figures 3 and 4)

**Figure 3.**
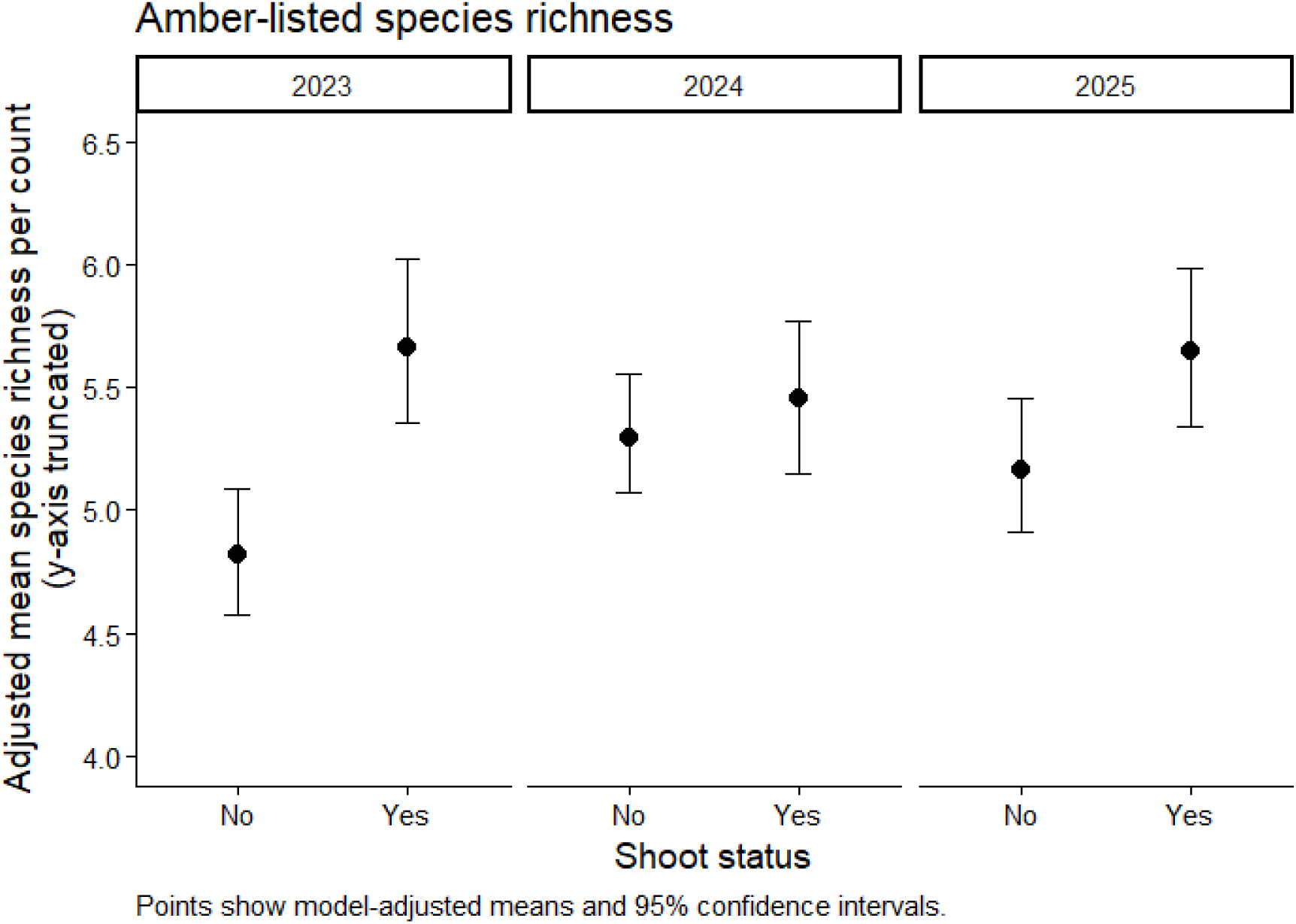
Model-adjusted mean Amber-listed species richness per count in shoot and non-shoot farms, 2023–2025. Points represent adjusted means and error bars indicate 95% confidence intervals.

**Figure 4.**
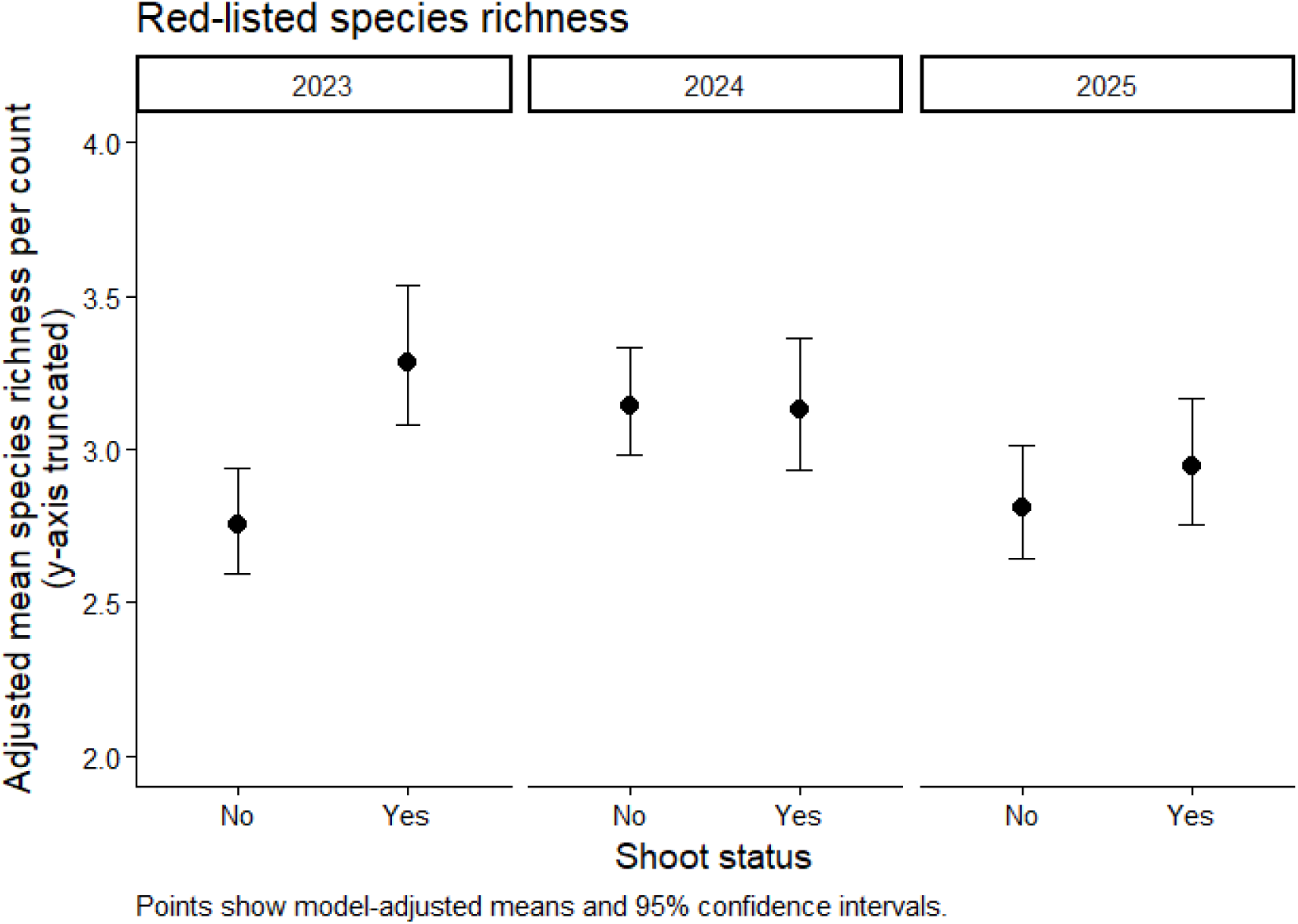
Model-adjusted mean Red-listed species richness per count in shoot and non-shoot farms, 2023–2025. Points represent adjusted means and error bars indicate 95% confidence intervals.

For the two groups of finches / buntings and thrushes, positive associations with shoot status were observed in 2023, with estimated species richness 28.9% (p < 0.001) and 14.3% (p < 0.01) higher on shoot farms respectively. Associations were smaller and not statistically significant for either group in 2024 or 2025. In contrast, raptor species richness showed statistically significant positive associations with shoot status across all three years, with estimated richness 20.7% higher in 2023 (p < 0.01), 23.0% higher in 2024 (p < 0.001), and 14.0% higher in 2025 (p = 0.03; Figures 5-7).

**Figure 5.**
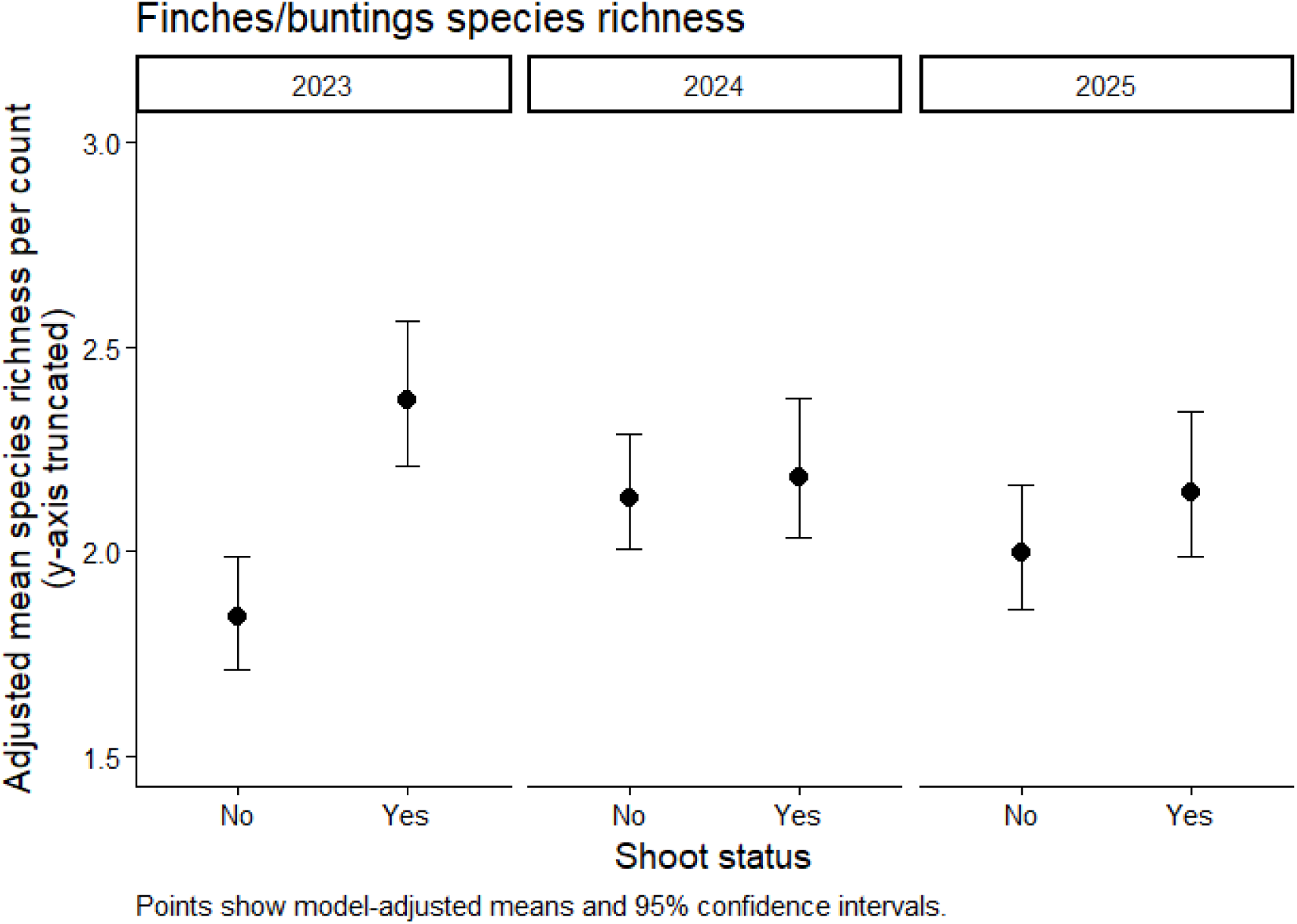
Model-adjusted mean finch/bunting species richness per count in shoot and non-shoot farms, 2023–2025. Points represent adjusted means and error bars indicate 95% confidence intervals.

**Figure 6.**
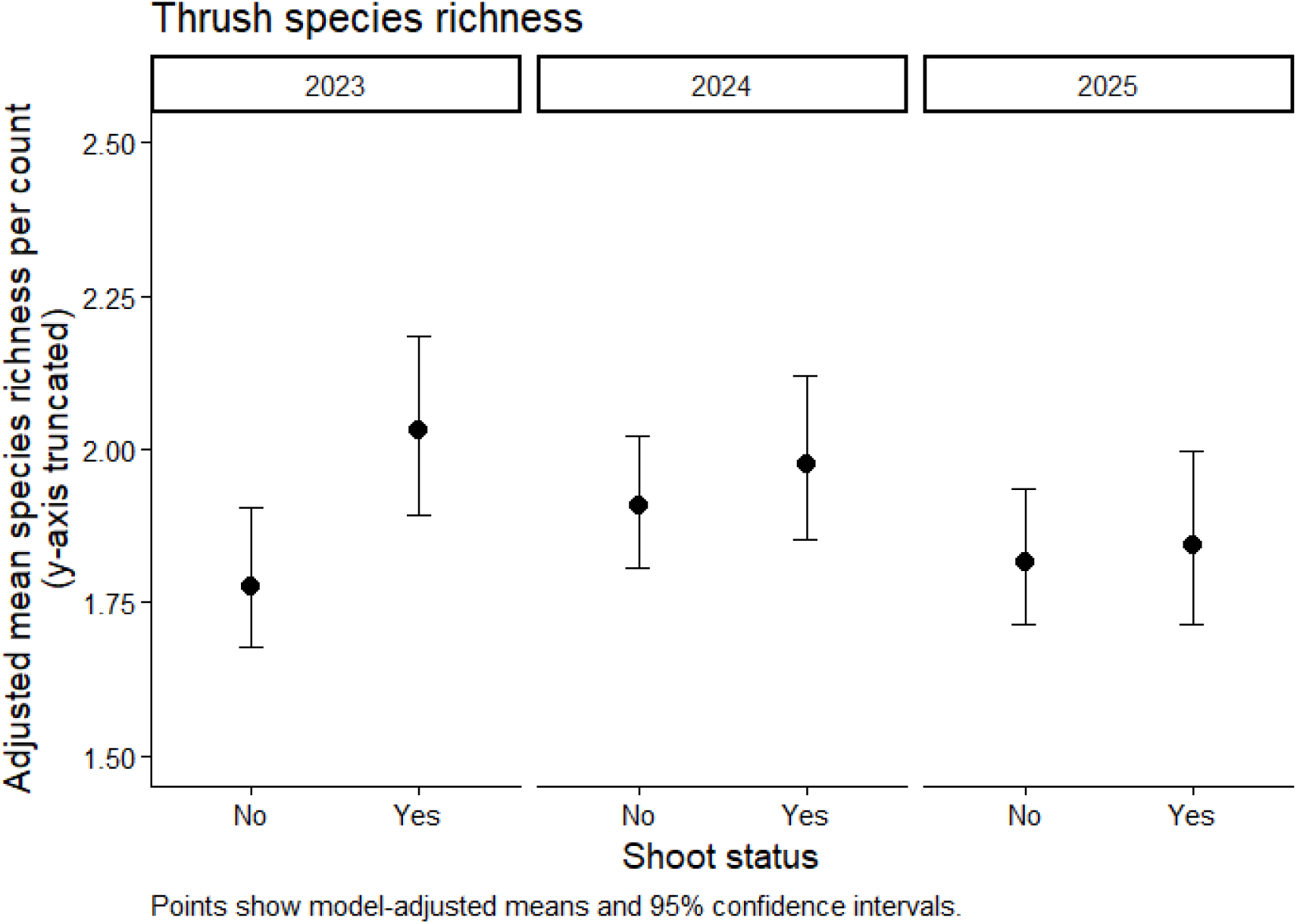
Model-adjusted mean thrush species richness per count in shoot and non-shoot farms, 2023–2025. Points represent adjusted means and error bars indicate 95% confidence intervals.

**Figure 7.**
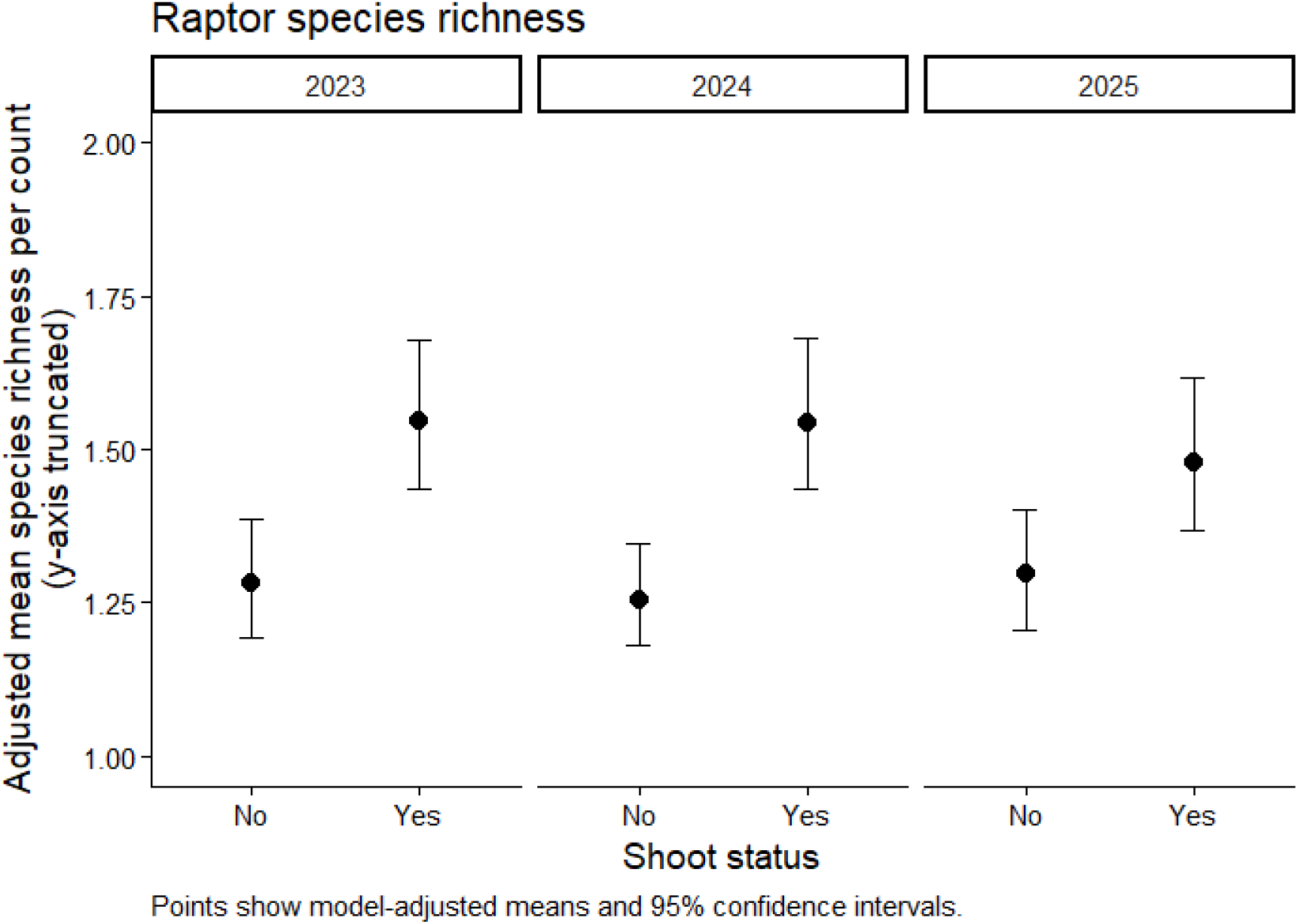
Model-adjusted mean raptor species richness per count in shoot and non-shoot farms, 2023–2025. Points represent adjusted means and error bars indicate 95% confidence intervals.

### Model diagnostics and sensitivity analyses

Model diagnostics generally indicated an adequate fit, with no significant evidence of residual dispersion or zero inflation. Specific sensitivity analyses restricting the FBI richness models to England only and excluding influential observations produced similar estimates to the main analyses, suggesting that these factors did not substantially influence the associations in the models.

## Discussion

The area of farms with shoots was on average substantially larger than that of non-shoot farms. Although surveys were undertaken on a standard 2ha patch of ground, this patch was not selected randomly and there is an increasing choice with larger areas. Shoots were also more likely to have AES options and less likely to be in a Farmer Cluster, both of which might also influence bird use. Our analysis removes the effect of farm size and of these other covariates to properly test whether the effect of being a shoot or not is still a significant factor for bird species richness.

While being a shoot appears to be a driver for AES option uptake, the effect of this is excluded from the analysis.

Using this strict approach, we found either higher or similar (i.e. no difference) bird species richness on the shoot sites for all species groupings in all three years investigated i.e. for none of the 21 group / year combinations was our measure of bird species richness higher, either significantly or otherwise, on the non-shoot group of sites. While we need to be careful with our interpretation, not least because as indicated in the methods our data are not fully independent between years, this overall result strongly suggests that there is a positive effect on bird species richness of gamebird management on farms with shoots.

Overall, there was significantly greater bird species richness on farms with shoots in 2023 and 2025. In 2024 there was no significant difference. The finches / buntings group would have contributed to these results as indicated. They are largely seed eaters and probably benefit from seed bearing game crops in winter and supplementary gamebird feeding into the spring (Henderson et al. 2003). There were also significantly more thrush species in 2023. Looking at indicators and listed species groups, there were more Amber listed species in 2023 and 2025 and more Red listed species in 2023 with no difference in other years. There were significantly more FBI species in 2023 and in 2025 there was a marginally insignificant difference (95% confidence interval: 0.99 to 1.15) for this group. Raptors as a group were 20% more common on areas with shoots in all three years. The commonest species encountered were red kites and buzzards, and these generalist predator species were probably responding to the presence of gamebird carrion in the winter and possibly young gamebirds in the autumn (Swan et al. 2022). There was a tendency for there to be numerically more kestrel as well although we are unable to test for this.

The survey farms are not a random sample, and the process of self-selection might involve some or other tendency. In particular, it is likely that participants in this survey would tend to be generally more interested in birds or wildlife, not least because there needs to be some ability to identify birds. However, there is no reason to suggest that participants ticking the ‘shoot’ box would be more interested in birds than participants not ticking that box. It is difficult to see how our main comparison between two types of sites might be distorted or biased by these processes, even if participants overall tend to be more interested in birds than average.

We are also aware that with three years data, the most appropriate analytical approach would be to combine data across years. The complication is that some sites contribute in more than one year, while others do not. To account for this, i.e that some data are not independent between years, we need to have a common site identifier in the dataset. This is currently unavailable and it is not possible to remedy that at this time. Results from each year are therefore to some extent not independent of each other in the analysis presented and we need to take this into account when interpreting them.

The bird data collected are of very variable quality with different count methods, timings and potentially bird ID accuracy. However, we do not expect these factors to cause bias in our comparison. Although participants do give counts of species, overall the data provide no consistent insight into abundance or density and so we have converted counts to site presence and absence. We suggest data in this form from a large sample of sites are suitable for looking at changes over time (which we are not considering here) as well as the relative effect of simple site factors such as shoot presence.

As indicated whether the bird survey area is part of a shoot or not is the only game management related question in the survey, so we do not know, for example, about the level of releasing or presence of game crops. Based on the GWCT National Gamebag Census survey over the last 10 years or so around 75% of lowland ‘shoots’ release pheasants and/or partridges (Aebischer 2019). On releasing shoots there may be negative effects of high densities of pheasants and/or red-legged partridge on other bird species. Wood edge and hedgerow habitats can be degraded where high densities of released pheasants occur (Sage et al. 2020).

There are possible zoonotic disease issues that might affect some birds (Madden & Sage 2020). It has been shown that where birds congregate, either at release pens or potentially at gamebird feeding sites, soil can be degraded and invertebrate communities altered or reduced. Away from release areas recent reviews suggest that these effects do not occur (Sage 2026).

Despite this potentially negative process, we found more or similar wild bird species on shoots in virtually all models tested in our data investigation. While bearing in mind the caveats outlined about data quality and the analysis, this suggests there is generally similar or greater bird species richness on farms with a shoot. If this is correct, it is appropriate to look in more detail at the mechanisms that might have caused the positive effect, including planting of game crops and provision of supplementary feeding, the maintenance of field margins and hedges, predator control, and possibly other conservation measures, all of which are commonly associated with the presence of a shoot (Sage et al. 2020).

Considering the first of these, the presence of seed-bearing game crops and associated supplementary feeding in winter and spring on 80% of shoots (Ewald 2004) is likely to be a factor in attracting wild birds to game managed areas of the farmed countryside. Winter game crops are widely planted on released game estates to provide feeding areas and suitable ‘holding’ areas for pheasants and red-legged partridge as part of a shoot (GCT 1994). Ewald (2004) also estimated that shoots planted on average 3% of the cropped area to game crops. A series of studies across the UK have shown similar results. Game crop plots or similar winter seed bearing crops typically have been found to contain around 10 or 12 songbirds per ha in most months between October and January, with the adjacent arable field plots typically containing less than one (South England: Sage et al. 2005; Scotland: Parish and Sotherton 2004; Midlands, Stoate et al. (2003), England-wide: Henderson et al. (2003).

In the early spring, after the shooting season, some game crops are annual and are quickly ploughed in, while some are biennial and others perennial. In most cases the provision of seed dries up in late winter (Henderson et al. 2003) so unless the gamebirds continue to be supplied with supplementary feed the food resource dries up. However, in a study looking at spring and early summer farmland bird populations using hedgerows around game crops and in places away from those crops, the effect of the game crops on birds was still apparent (Sage et al. 2024).

They found that otherwise similar hedges in terms of size and density near to those winter game crop plots, had between 1.5 and 2 times as many breeding resident songbirds per unit length the following spring compared to hedges further than 350 m away from winter game crop plots.

Looking at a second factor, it is likely that many hedgerows today, like woodlands, were planted or retained for game interests in the past (Rackham 2003). Hedgerows are used by game managers to help connect woodland pheasant releasing areas to game crops, and to provide habitat for red-legged partridges in open field release areas, to facilitate shooting (GCT 1994; 1996). Firbank (1999) reported more hedges, more complete hedgerow networks and greater connectivity between hedges and woods on game areas than on non-game areas. Using satellite imagery Draycott et al. (2012) found that 97 release-based pheasant shoots had 10% to 65% more hedgerow per square kilometre than 53 farms with no releasing. Using ground surveys, while the size of the hedges on game and non-game sites were similar, game hedges were more likely to have a grass margin or other uncultivated strip alongside it. In an assessment of forest and hedgerow metrics Long et al. (2025) found greater structural diversity and hedgerow density across 1100 or so known pheasant-managed areas and 1100 matched control areas. This result was apparent despite the fact the study did not verify whether control sites had game managed activities or not.

Many farmland bird species need hedgerows to nest and or forage in (Hinsley & Bellamy 2000). Some of the hedge characteristics relating to size, density and adjacent habitats that these studies indicate are more common on game managed areas have also been shown to be better for birds (Arnold 1983; Green et al. 1994). Draycott (2012) recorded more birds in game hedges and Sage et al. (2024) documented 1.5 to 2 times more birds in hedges within 350m of game crop plots than hedges further from those plots.

The third main factor that may be relevant to our findings, i.e. farms identifying as shoots in our study sometimes had increased species richness, is predator control. In a wide-ranging review Reynolds & Tapper (1996) describe and discuss how predator control to benefit small game has been practiced in the UK for 200 years and has played a key role in shaping the UK fauna today. A theme in this area of research is that where predator control is undertaken effectively alongside beneficial habitat management, then birds and other wildlife can benefit. For example, White et al. (2014) showed that this combination had a measurable benefit on nest success for several hedgerow nesting bird species at a game managed site over many years. By comparing the effects of management on avian abundance at the GWCT and RSPB demonstration farms over 25 and 15 years respectively, Aebischer et al. (2016) concluded that where predator densities were high (>3 corvids and 1 fox per km^2^), it is likely recovery of declining farmland birds including some passerines, would require predator control as well as habitat management.

In their wide-ranging review of UK predation of wildlife Roos et al. (2018) provided good evidence that for the main bird groupings with available evidence (seabirds, gamebirds, waders, and to an extent, passerines) predation can sometimes be limiting and that predator management aimed at foxes and corvids simultaneously (as applied by many lowland shoots) can be effective. In their review Mustin et al. (2018) concluded that predator control associated with game management had a positive effect on some wildlife.

It has been thoroughly demonstrated that fox control by gamekeepers can reduce fox populations locally or regionally (Reynolds et al. 1993; Tapper et al. 1996; Porteus, 2015). There is also some evidence that coordinated collaborative fox control can result in reducing the population of foxes regionally. (Heydon & Reynolds (2000) documented this in one of three UK regions studied). The missing link here is the extent to which modern lowland shoots, most of which are based on releasing pheasants and partridges, undertake effective fox and corvid control. The studies outlined here and others reviewed in Roos et al. (2018) suggest that some do. In their review, Sage et al. (2021) concluded that there is some evidence to suggest that probably a minority of release-based shoots, undertake effective predator control. In these circumstances the studies outlined here and others reviewed in Roos et al. (2018) suggest that birds on farmland and in other habitats can benefit in terms of productivity metrics and possibly adult survival.

A fourth factor is the possibility that game managed shoots do more habitat management and/or other conservation work, that is not directly related to the management of game. This possibility is supported by little evidence. However, a game manager or gamekeeper is generally someone working on an area of land and not entirely distracted by crop or livestock productivity. It is reasonable to suggest that such a person will often be considering habitat quality as well and might have more scope to do some conservation orientated work, than say the farm manager or a tractor driver.

In summary it is likely that these activities may work in combination on most releasing and some other gamebird shoots in the UK to enhance the game managed area so that more birds use those areas throughout the year.

Sharps et al. (2023) provides an insight into the scale of the requirement in terms of higher and lower tier AES uptake, to reverse the decline in farmland birds (the FBI) from the current situation in terms of habitat provision. They concluded that to reverse overall declines by 10% over 10 years a combination of higher and lower tier AES together with targeted wild bird options, 30% of arable land and 10% of pastoral land would be needed.

To make a basic comparison, we can derive an estimate of how much UK lowland farmland might be benefitting from game management associated with the presence of a shoot. Ewald (2004) estimated 80% of all shoots planted game crops covering 3% of their arable area and this tallies with unpublished material about the deployment of game crops. Madden et al. (2021) uses a variety of data sources and approaches to make estimates of how many lowland pheasant / partridge shoots employing at least one part-time gamekeeper (who can do some management work) there are in the UK. The median figure is 7000, and the paper suggests this is realistic.

There are several online based surveys of shoots in the UK, the largest of which is run by Guns on Pegs and Lycetts (GOP 2025). Amongst other things participants are asked about shoot area. A typical figure from these surveys in recent years is that the average shoot area from a sample of between 500 and 1000 shoots is about 600ha. In the BFBC survey the farm area for the shoot group was about half that i.e. 300ha. Which of these provides the most representative figure is not clear so for our broad comparison we will choose an average of 450ha. Combining this figure with the 7000 shoots figure and discounting the 20% of shoots that don’t plant game crops, means that 24 million ha or 24 000 km2 is to some extent managed as a shoot with game crops.

With a total UK lowland farmland area of 125 000 km2 (ONS 2022) we arrive at an estimate that about 20% of that lowland farmland contains a shoot that plants game crops. This falls midway between the figures estimated for arable and grass by Sharps et al. (2023) in relation to AES options and reversing FBI declines.

Our study shows that over the three years of data we have, the FBI species group counts were on average about 10% higher on shoots than on non-shoots. This excludes the effect of AES options, which were more commonly deployed on shoots. If this is correct, on these places the FBI index declines would have been noticeably less steep over the last 20 years. Taking account of the proportion of managed shoot area figures above across UK lowland farmland, we estimate that if game management stopped, the national overall FBI index figure today would be 2% lower than it actually is.

We cannot say that game management on an average shoot is more or less useful than a higher tier AES agreement in terms of its benefit to birds. But given the scale of game management and the measurable benefit it has to many farmland birds indicated by this investigation, it is reasonable to suggest the requirement in terms of costly stewardship options to maintain the FBI where it is now without gamebird management, would be higher than it currently is.

